# Use of unmanned aerial vehicles (UAVs) for mark-resight nesting population estimation of adult female green sea turtles at Raine Island

**DOI:** 10.1101/2020.01.21.913681

**Authors:** Andrew Dunstan, Katherine Robertson, Richard Fitzpatrick, Jeff Pickford, Justin Meager

## Abstract

Nester abundance is a key measure of the performance of the world’s largest green turtle rookery at Raine Island, Australia. Abundance surveys have been undertaken in waters adjacent to Raine Island reef using mark-resight counts by surface observer (SO), underwater video (UWV) and unmanned aerial vehicle (UAV) (since 1984, 2013 and 2016 respectively). UAV and UWV may provide more cost-effective and less biased alternatives, but estimates must be comparable with the historical estimates. Here we compare the three methods.

The relative likelihood of resighting a marked turtle was significantly higher by SO than the other methods, which led to lower mark-resight population estimates than by UAV or UWV. Most (96%) variation in resighting probabilities was associated with survey period, with comparatively little variation between consecutive days of sampling or time of day. This resulted in preliminary correction factors of 1.53 and 1.73 from SO-UWV and SO-UAV, respectively. However, the SO and UWV estimates were the most similar when turtle densities were the lowest, suggesting that correction factors need to take into account turtle density and that more data are required.

We hypothesise that the UAV and UWV methods improved detection rates of marked turtles because they allowed subsequent review and frame-by-frame analysis, thus reducing observer search error. UAVs were the most efficient in terms of survey time, personnel commitment and weather tolerance compared to the SO and UWV methods.

This study indicates that using UAVs for in-water mark-resight turtle abundance estimation is an efficient and accurate method that can provide an accurate adjustment for historical abundance estimates. Underwater video may continue to be useful as a backup alternative to UAV surveys.

## Introduction

Green turtles, *Chelonia mydas*, are listed as vulnerable in the State of Queensland (Nature Conservation Act 1992) and in Australia (Environment Protection and Biodiversity Conservation Act 1999). The majority of the northern Great Barrier Reef (nGBR) population of green turtles nest at Raine Island (Fig 1), which is the world’s largest remaining green turtle rookery (Seminoff et al 2015). Concerns about low reproductive success of green turtles at Raine Island have been reported since 1996 (Limpus et al 2003; Dunstan et al 2018), which is thought to caused by nesting beach inundation as well as nest environment factors such as respiratory gas, microbial or temperature extremes (Dunstan et al 2018). The population is also exposed to other cumulative impacts including climate change (Fuentes et al 2011), feminisation (Jensen et al 2018), hunting (Graysona et al 2010), plastic pollution (Schuyler et al 2014), vessel strikes (Hazel et al 2006), commercial fishing (Wilcox et al 2015) and coastal development (Bell et al 2019). An accurate index of nesting population numbers is critical for understanding the reproductive success and long-term changes to population numbers.

**Fig 1.**
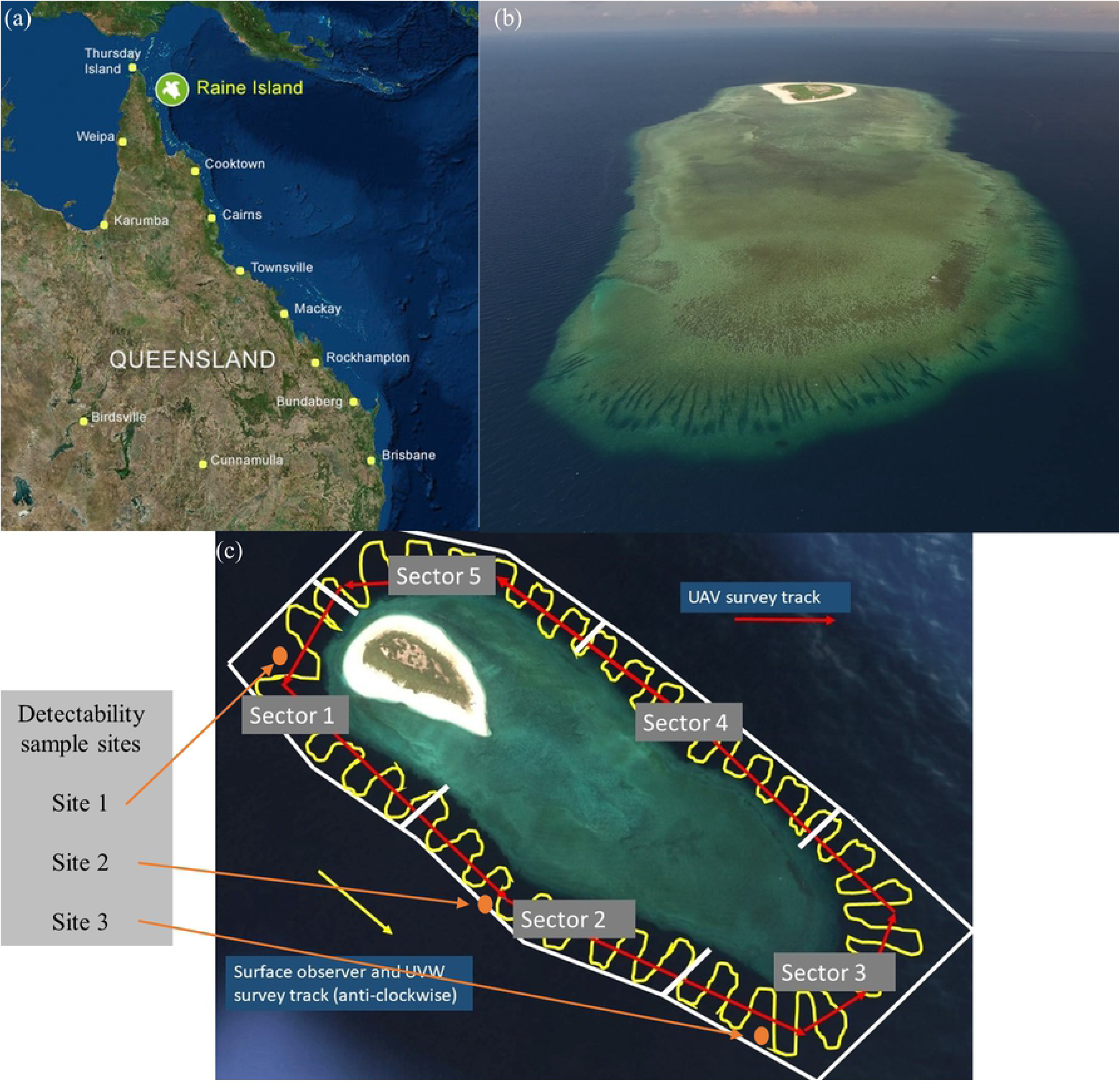
Raine Island location. (a) Location of Raine Island on the northern Great Barrier Reef, Australia, (b) Raine Island reef study site and (c) transect search paths with turtle detectability experimental sample sites marked.

The remoteness of Raine Island and the sheer number of nesters on a given night have precluded total nesting censuses or a comprehensive mark recapture program. Instead, a mark-resight approach has been used to estimate the numbers of nesters in the surrounding internesting habitat since 1984 (Limpus et al 2003). Females are painted (marked) during nightly tally counts, and counts of marked and unmarked turtles in the waters that surround Raine Island are used to estimate abundance during the sampling period using the Lincoln-Petersen estimator (LP). Mark-resight is therefore combined with in-water sampling, and thus estimations of nester abundance are dependent on the limitations and assumptions of both approaches.

The major challenge for in-water surveys is to have high detectability for both marked and unmarked turtles, given that marine turtles spend only a small proportion of their time at the water surface, especially when surface conditions are poor, in turbid water or when turtles are amongst habitat structure (Fuentes et al 2015). The LP estimator is based on the assumption that the population is ‘closed’ during the sampling period (Williams et al 2002), which means that they do not depart from inter-nesting habitat in the short time interval from marking to the in-water survey. Another key assumption of the LP estimator is that the probability of detection is the same between marked and unmarked turtles. The LP estimator is also only based on one resighting event, which could make it less robust than estimates from repeated sampling.

The introduction of modern technologies such as UAVs and underwater video for counting surveys coupled with artificial intelligence for automated image analysis may provide a more time efficient and reliable mark-resight estimate. Another advantage of UAVs and underwater cameras compared to the vessel platform is that effect of surface reflections can be supressed or eliminated. Here we aimed to compare the effectiveness of the vessel observers, UAVs and underwater video, and to determine if the UAV and underwater camera estimates are comparable to the historical data. Our comparison of detectability of marked turtles between methods also provided the opportunity to test the key LP estimator assumption of equal detectability of marked and unmarked turtles. If the probability of detection is the same between marked and unmarked turtles, the ratio of marked and unmarked turtles should not differ between sampling methods. Finally, we used a repeated sampling study design to (a) determine whether there is a gain in precision in the LP estimator with repeated sampling, and (b) test whether the closure assumption was appropriate.

## Materials and methods

### Ethics statement

All procedures used in this project were approved by the Raine Island Scientific Advisory Group and by the Queensland Department of Agriculture and Forestry Animal Ethics Committee (Permits SA 2015/12/533 and SA 2018/11/660).

### Study area

Raine Island is located on the outer edge of the northern Great Barrier Reef and is part of the Raine Island National Park (Scientific). The Wuthathi People and Kemerkemer Meriam Nation (Ugar, Mer, Erub) People are the Traditional Owners and Native Title holders for this country and are an integral partner of the area’s management. Over thousands of years, Wuthathi People and Kemerkemer Meriam Nation People have held cultural connections to Raine Island through the use of its resources and cultural connections to the land and sea, through song lines, stories, and voyages to the island.

All research was undertaken on the reef waters adjacent to the Raine Island National Park (Scientific) (11° 35’ 25” S, 144° 02’ 05” E) between November and February during the 2013-14 to 2017-18 green turtle nesting seasons (Fig 1). Raine Island reef has a perimeter of approximately 6.5 km and is fringed by coral reefs. Green turtles are the only sea turtle species recorded nesting at Raine Island where the nesting beach is approximately 80 m wide with a circumference of 1.8 km. Nesting is seasonal with the main nesting period from October to April and extremely low rates of nesting for the rest of the year (Limpus 2007). The peak nesting period is from December to January.

As many as 23,000 turtles have been counted in one night at the beach. However, there is a large variability in green turtle nesting numbers from year-to-year that is correlated with the lagged Southern Oscillation Index (Limpus and Nichols, 2000).

### Turtle marking procedure

The carapaces of nesting turtles were painted along the midline with a white stripe approximately 80 cm in length and 20 cm in width, using a 12 cm wide paint roller and “APCO-SDS fast dry water-based road marking paint” (MSDS Infosafe No. 1WDKY) (Dunstan, 2018). A turtle was selected for painting if the carapace was dry, the carapace did not have a thick coating of algae and the turtle was inland of the beach crest (to provide sufficient time for the paint to dry). When applied under these conditions, the paint adhered to the carapace surface for at least 96 hr. This was confirmed over the three nights following painting of turtles, which provided the opportunity to assess the paint when many painted turtles came ashore to re-attempt nesting. While there was erosion of paint on a small proportion of turtles, enough paint always remained to allow identification of turtles as ‘painted’.

Turtles were painted on a single night during turtle survey trips in November (2016), December (2013, 2014, 2016, 2017) and February (2016). All suitable turtles on the nesting beach were painted, up to a maximum of 2000 (Table 1). The upper limit was determined by logistical constraints and time while the lower limit was influenced by nesting turtle numbers.

**Table 1.** Summary of survey periods, number of turtles marked and survey methods conducted.

| Survey period | Marked turtles | Number of surveys |  |  |
| --- | --- | --- | --- | --- |
|  |  | Vessel observer | UVW | UAV |
| Dec 2013 | 2000 | 6 | 1 | - |
| Dec 2014 | 1930 | 3 | 3 | - |
| Feb 2016 | 482 | 5 | 6 | - |
| Nov 2016 | 781 | 6 | 6 | - |
| Dec 2016 | 2000 | 6 | 5 | 3 |
| Dec 2017 | 2000 | 2 | 3 | 3 |

### In-water detectability of marked and unmarked turtles

We tested the detectability of submerged green turtles using a model, which was constructed from plywood to represent an average-sized nester with curved carapace length of 106 cm (Limpus, 2003) and painted appropriately. The model was lowered on a rope and the depth at which it was no longer discernible as a turtle was recorded. A painted white plywood board the same size and colour as the turtle marks was then attached to the model to simulate a marked turtle. The model was again lowered to determine the depth that the white marking was still obvious. Single samples for each treatment were undertaken at three locations that represented the range of water conditions around the island from coastal aspect (site 1) to between reef channel (site 2) to open ocean aspect (site 3) (Fig 1c).

### Mark-resight counting methods

Surveys were undertaken between November and February during 2013 to 2017. Turtles were counted if the turtle shape was discernible and the presence/absence of the painted white mark was recorded. A pilot study using SO, UWV and UAV methods indicated that the white markings were visibly obvious and the presence-absence of the mark was never in doubt. All unmarked turtles were considered to be adult female turtles, because previous surveys (Limpus, 2003) demonstrated the minimal presence of adult males and juveniles during the survey period. Wind speed was mostly low during the surveys (average maximum wind speed: 11 knots, range: 1 to 18.7 knots). Water clarity measured at three sites around Raine Island using a standard Secchi disc (30 cm diameter) ranged from 9 to 13 metres.

#### Surface observer method (SO)

A standardised search area was surveyed in the waters surrounding the island on the morning and afternoon of the three days following turtle marking, or less where logistics limited sampling (Table 1 & Fig 1c). A 4.2 m outboard powered rigid hull inflatable vessel with three persons aboard, one recording, one driving and one counting, was driven along the waters adjacent to the reef perimeter edge in search of the painted turtles (Fig 1c).

#### Underwater video method (UWV)

Underwater video surveys were conducted from the survey vessel simultaneously with the surface observer surveys (Table 1 and Fig 1c & 2a). A GoPro Hero4 camera (frame rate: 25 hz; resolution: 1080; field of view: 127°) with an extended life battery was attached to the hull of the vessel pointing forward and downward, and recorded throughout the entire reef perimeter survey period. Video footage was reviewed by one observer using a single tally counter to record female turtles that could be scored positively for turtle shape outline and for the presence/absence of the white paint mark during separate video replays. Video playback was paused during peak turtle density periods and playback speed adjusted for counting efficiency and accuracy.

**Figure 2.**
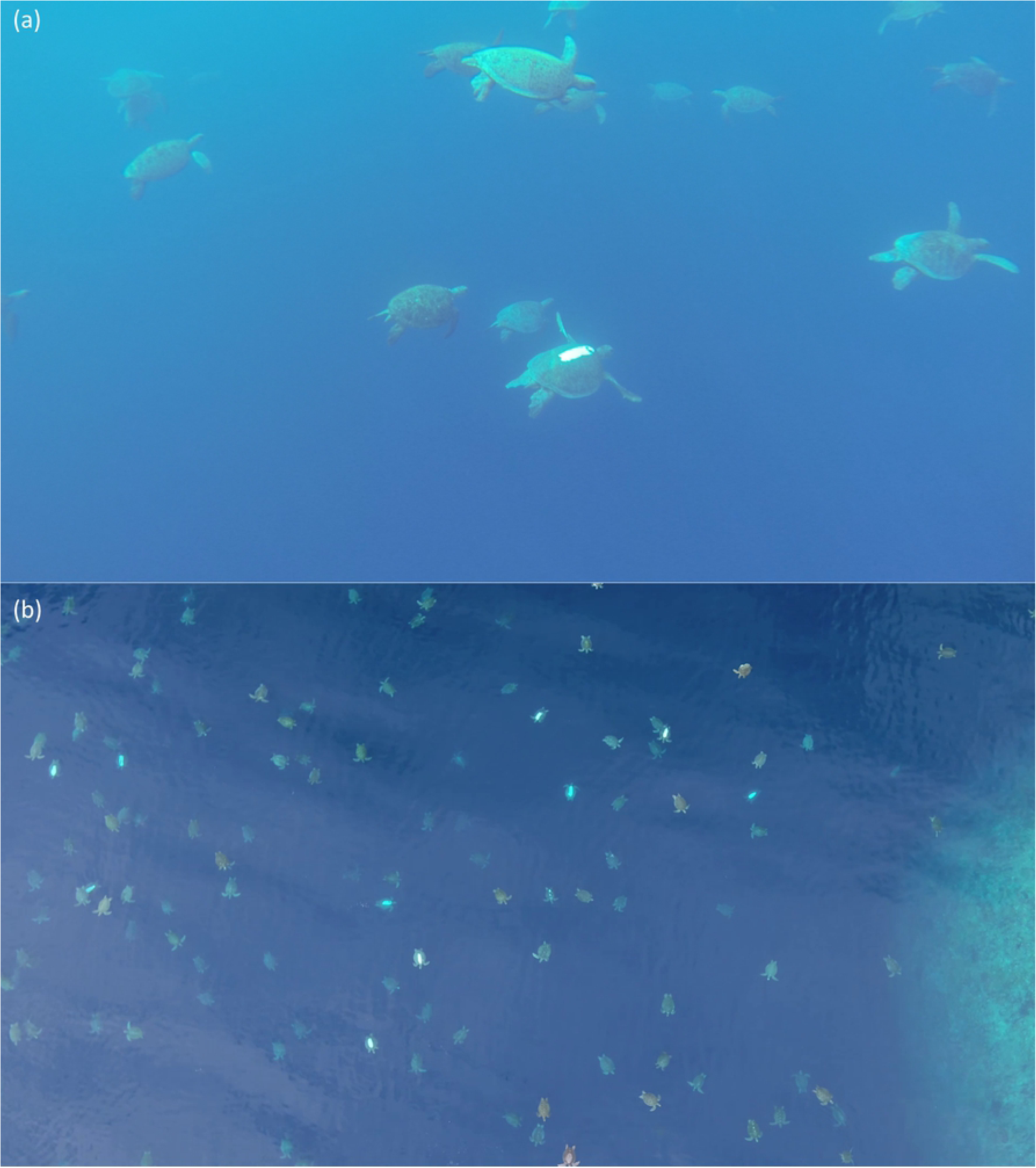
UWV and UAV survey image examples. (a) Still image from underwater video December 2017 survey and (b) still image from UAV video survey December 2017 at 50 m survey altitude.

#### UAV method

UAV surveys were conducted as close to midday as possible to reduce sun glare on the water surface. A DJI Inspire 1 UAV with Zenmuse X3 camera (frame rate: 25 hz; resolution: 1080; field of view: 94°) was flown at an altitude of 50 m and a speed of 5 m/s along a path consistent with that of the surface observer and underwater video surveys (Figs 1c & 2b and S1 multimedia). This camera and 20 mm equivalent lens provided a horizontal video survey swathe of 90 m at the sea surface. The UAV pilot was in the same vessel used for the surface observer and underwater video surveys, which followed closely behind the UAV. Video footage was analysed as described for UWV surveys.

### Statistical analyses

We first compared detection depths of the turtle model (with and without the painted mark) at the three sites using a Student *t* test on log_e_ transformed Secchi depths. We then compared the relative probability of detecting a marked (painted carapace) turtle between survey methods using a generalised linear mixed effects model (GLMM) with a binomial link function. A mixed-effects design was required because each batch of marked turtles was observed twice daily for five days. The optimal variance structure for the random effects was first explored using the ‘lme4’ package (lme4 v. 0.999375-35) of the R statistical environment (v. 2.13.1; R Development), using residual diagnostics and Akaike’s Information Criterion (AIC) of different mixed models (following Zuur et al., 2009). A model that allowed the slope of the day within nesting season effect to vary resulted in only a marginal improvement in AIC over a model that included a nested random effect of diel period (morning or afternoon) within day and nesting season. The relative probability (P) of detecting a painted green turtle (M) was therefore modelled by:

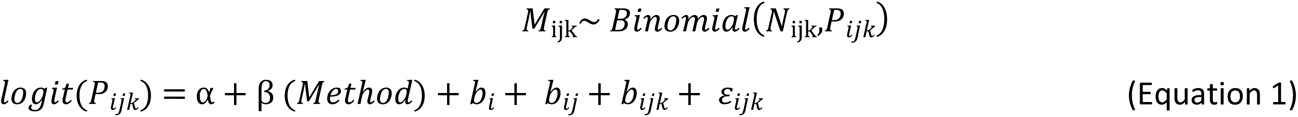

where the relative probability of detecting a marked green turtle (*P*) in a given time period (*i*), day (*j*) and nesting season (*k*) is a function of the survey method (*Method*). Other terms in the model are the total number of turtles that were resighted (*N*), the general intercept (α), the random intercepts (*b*) and the residual error (*ε_ijk_*). Equation 1 was fitted in a Bayesian framework using the ‘mcmcGLMM’ package and vague priors.

We then explored the gain in accuracy and precision in the LP estimator (Williams et al., 2002) from repeated recapture periods using a jacknife procedure. Each jacknife resample calculated population size as a function of the cumulative average of marked and unmarked recaptures, up to a maximum of six samples by the end of the third day (i.e. samples were taken twice daily for three days).

We estimated conversion factors for the historical estimates as the quotient of the mean population estimates, e.g. the conversion factor for SO to UWV estimate was the SO population estimate divided by the UWV population estimate. To explore how this conversion factor varied with population size, we fitted a linear regression of conversion factor against SO population size. Finally, we compared the number and densities of turtles sighted in each method using general linear models.

## Results and discussion

### In-water detectability of marked and unmarked turtles

The white mark was discernible at an average of 3 metres deeper than the turtle model (*t* = 3.61, df = 3.8, *p* = 0.026).

### Comparison of detectability between methods

Results consistently demonstrated a higher detection ratio of marked:unmarked turtles using UAV and UWV when compared with the SO method. Analysis of this data translates to significantly higher LP population estimates from the UAV and UWV methods compared to the SO method (Table 2 & Fig 3).

**Table 2.** Mean values for total mature female turtles counted and Lincoln Peterson estimates for periods surveyed by each method with standard error.

| Survey period | Vessel surface observer |  |  | UWV |  |  | UAV |  |  |
| --- | --- | --- | --- | --- | --- | --- | --- | --- | --- |
|  | Total turtles | Peterson estimate | S.E | Total turtles | Peterson estimate | S.E | Total turtles | Peterson estimate | S.E |
| Dec 2013 | 3167.2 | **58817.8** | 6095.1 | 4289.0 | **102142.9** | 10969.9 | - | - | - |
| Dec 2014 | 1002.7 | **14439.1** | 1174.1 | 534.0 | **18827.3** | 2470.9 | - | - | - |
| Feb 2016 | 169.2 | **4708.7** | 1116.3 | 194.8 | **5398.2** | 1351.5 | - | - | - |
| Nov 2016 | 728.8 | **8838.1** | 1074.4 | 1000.5 | **13180.8** | 1756.6 | - | - | - |
| Dec 2016 | 705.5 | **12377.5** | 1089.1 | 1275.8 | **18135.9** | 1496.1 | 1460.0 | **19682.9** | 1553.2 |
| Dec 2017 | 1596.5 | **20009.4** | 1618.7 | 1679.3 | **33263.1** | 3198.1 | 4622.3 | **37035.0** | 2334.0 |

**Fig 3.**
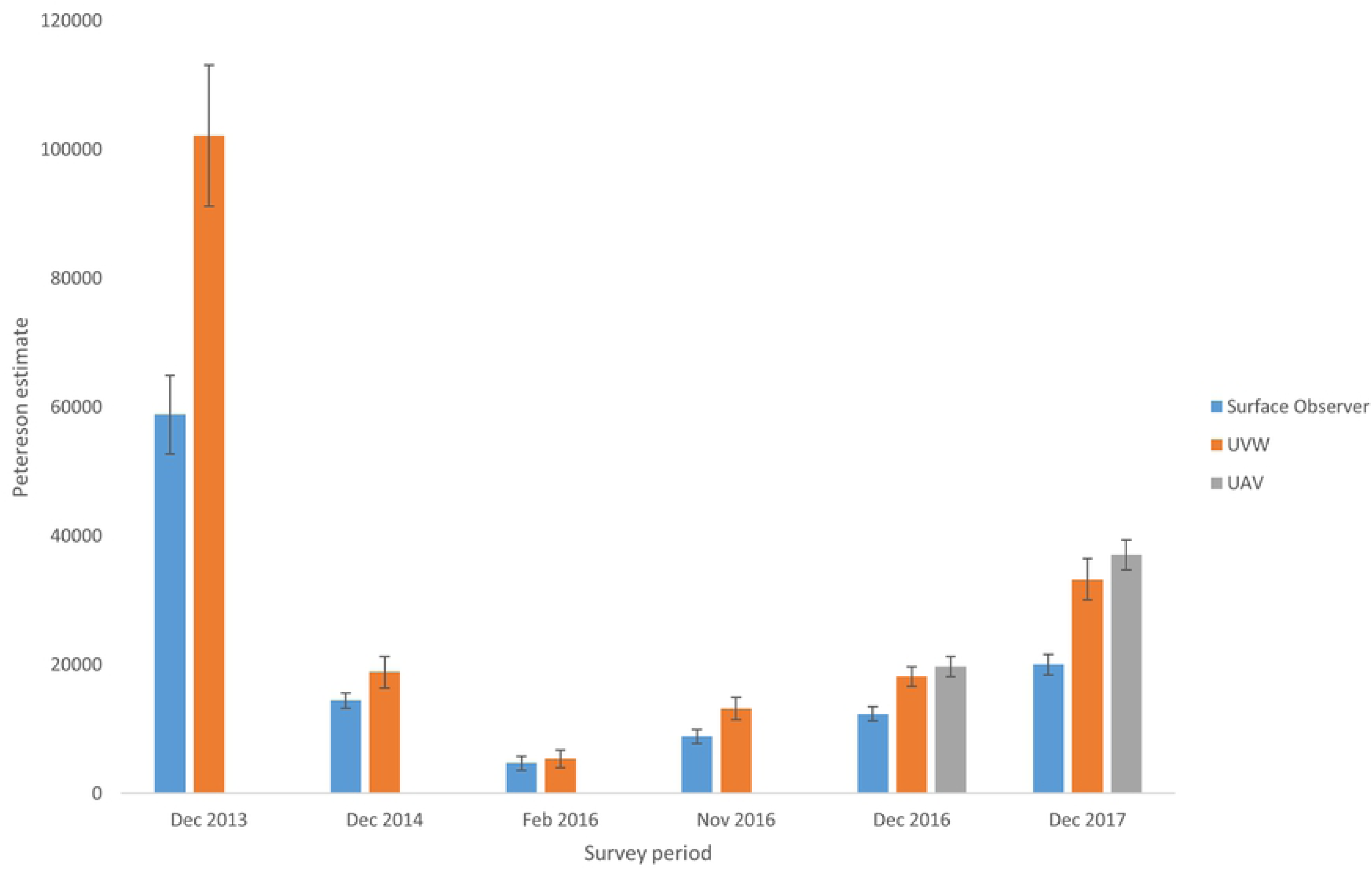
Lincoln Peterson population estimates for periods surveyed by each method. Error bars shown are ± 1 standard error.

Survey period accounted for 96.8% of variation (highest posterior density intervals from 82.6 and 99.6%) in the relative probability of detecting a marked turtle, compared to negligible variance components associated with sampling day (2.58 x 10-5 %, nested within sampling period) or time of day (5.17 x 10-5 %, nested within sampling day and sampling period). On average, 9.45 % of turtles detected using the SO method were marked (95% CI: 5.24% to 15.29%), compared to 6.58% for the UWV method (95% CI: 3.21% to 12.02%) and 6.26% for the UAV method (95% CI: 2.86 to 12.07%) (Fig 4).

**Fig 4:**
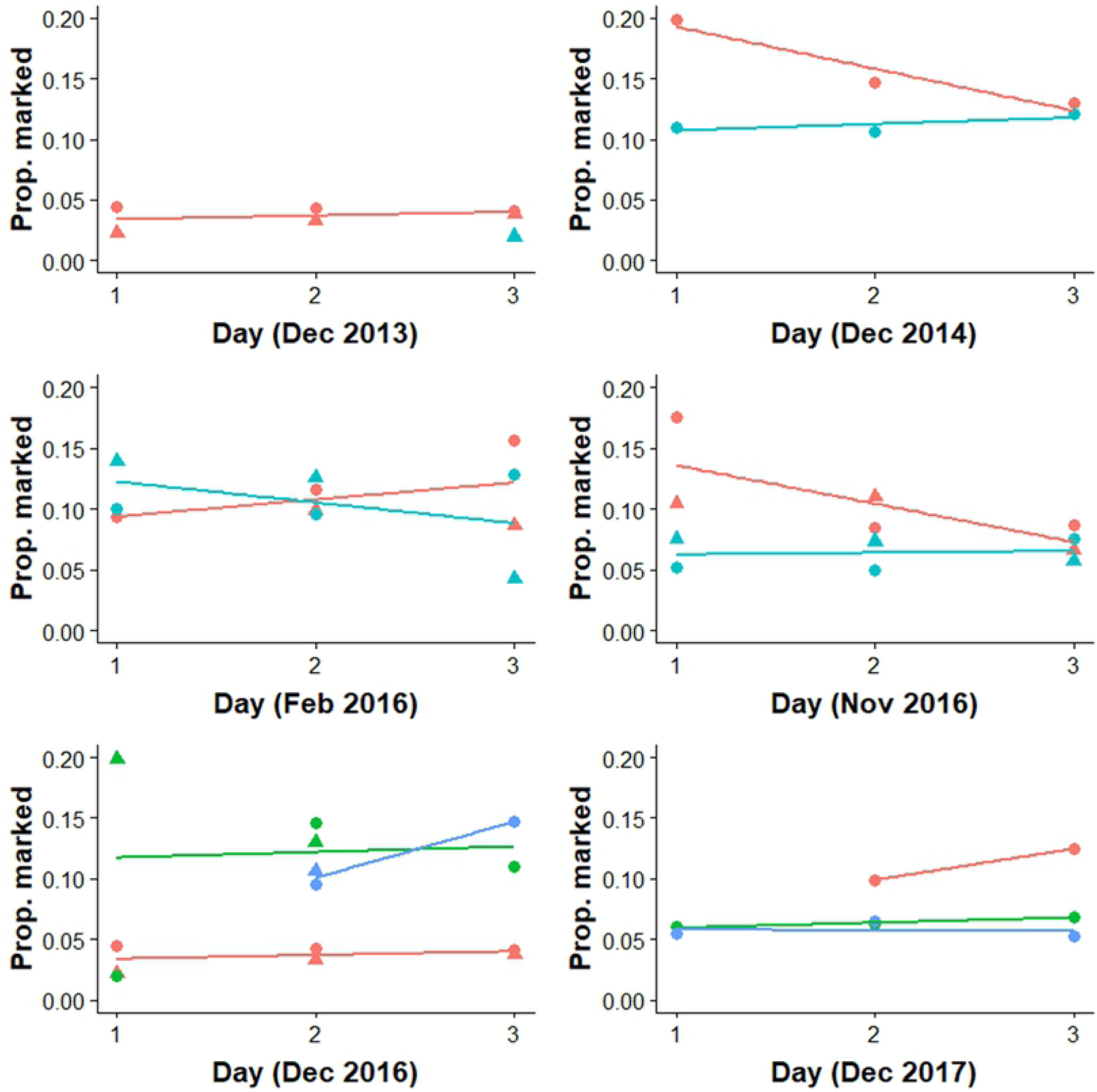
proportion of marked turtles detected for each method. Plot legend is (red: surface observer; blue, underwater video; green; UAV) and diel period (circles: morning; triangles: afternoon). Samples were collected over three successive days on each occasion. The coloured lines represent the average trend over time for each method and period.

Once variation associated with survey period was accounted for, there was no significant difference in detectability between the UAV and UWV methods (Fig 5).

**Fig 5:**
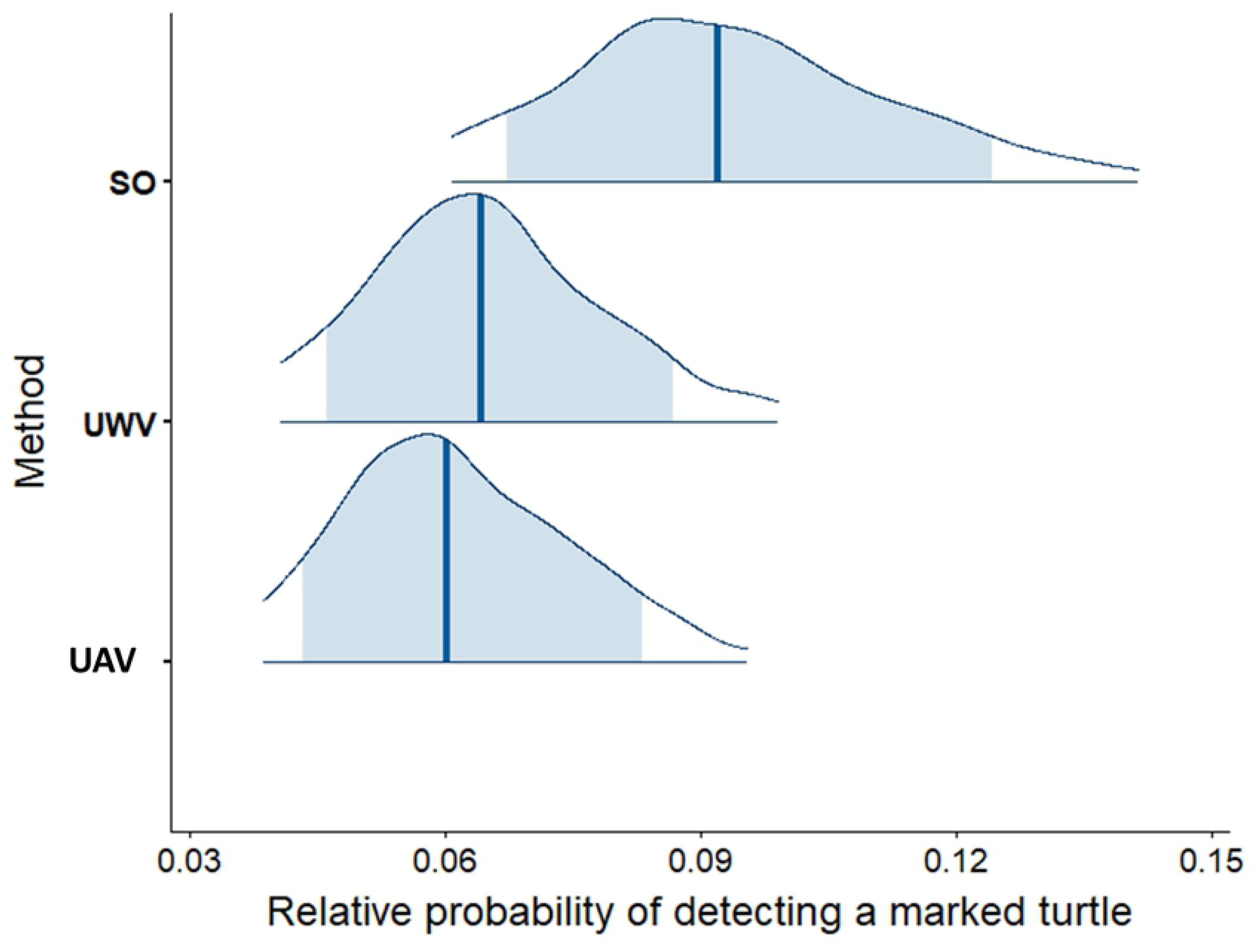
modelled relative probabilities of detecting marked turtles using each method. (SO, surface observer; UWV, underwater video and UAV). The density plots are computed from the merged posterior draws, where the blue vertical line represents the median and shaded blue areas are the 80% credibility intervals.

The relative gain in precision from using repeated measurements was similar across all three survey methods (Fig 6). There was an obvious gain in using two measurements (rather than one). Estimates and variances stabilise after three measurements suggesting that three measurements is sufficient.

**Fig 6:**
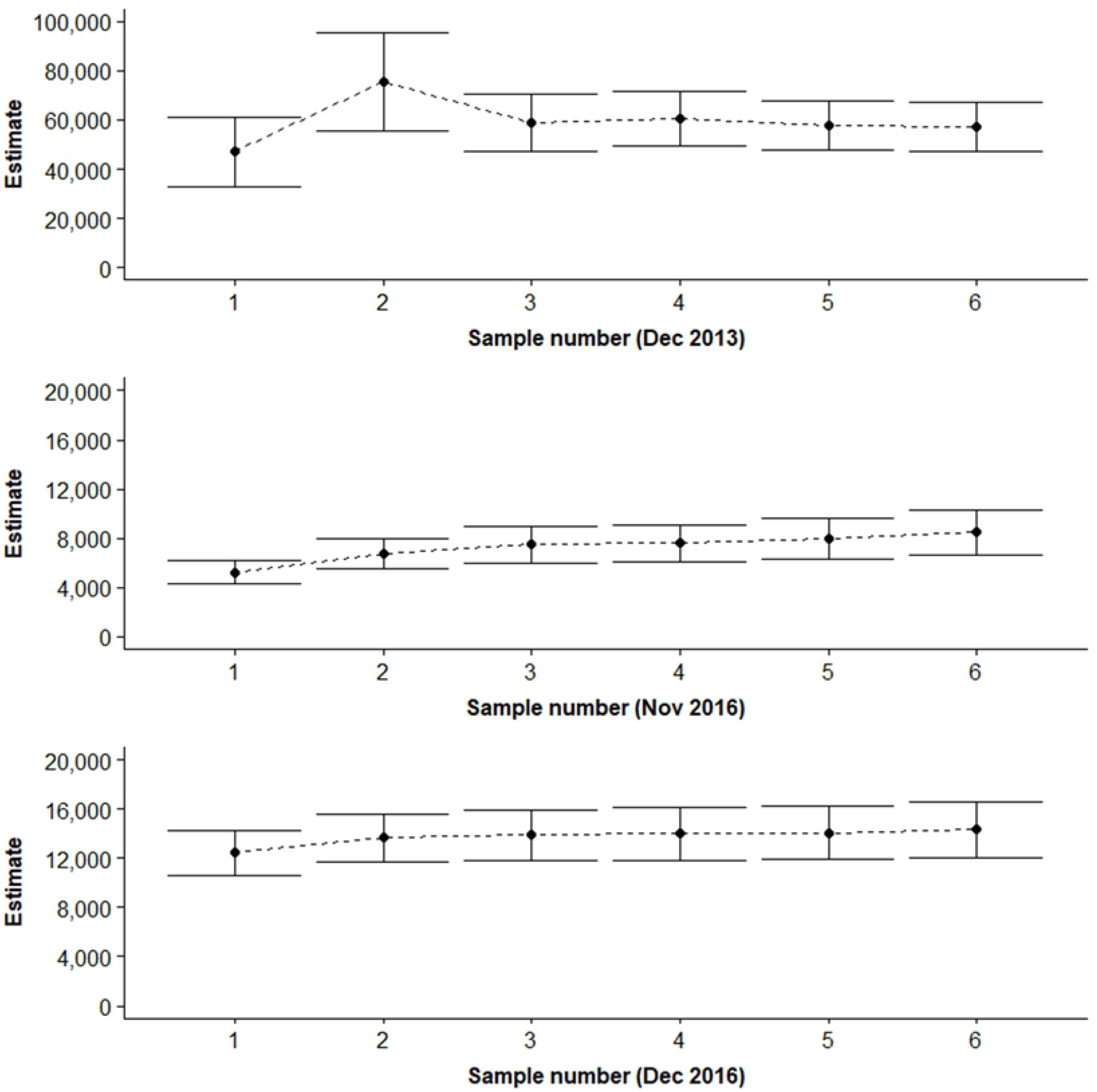
influence of sample size on the Lincoln-Petersen estimate. Shown here are the estimates for the Surface observer method and the three sampling periods for which six samples were available (± 95% confidence intervals).

Due to the major differences between Lincoln-Peterson estimates using Surface observer and both UWV and UAV techniques, the consistency and subsequent application of conversion factors was investigated. Conversion factors were calculated by dividing mean LP estimates between SO, UWV and UAV techniques where surveys using these techniques were conducted during the same time period. These conversion factors were then averaged to provide a mean conversion factor (SO to UWV CF = 1.53 (SD = 0.24), SO to UAV CF = 1.73 (SD = 0.18) and UWV to UAV CF = 1.11 (SD = 0.01)).

However, there was considerable variation in detection probabilities between sampling periods, which was likely to be driven by the extreme variability in the density of turtles in the inter-nesting habitat. Conversion factor calculations for UWV vs SO methods were compared with population estimates from SO surveys from different seasons. The results showed a significant linear relationship with a fitted regression line y = 1.3436e^6E-06x^ with an *r^2^* = 0.577 between UWV:SO conversion factor ratio and the seasonal nesting population density (Fig 7). F-test results show F < F Critical one-tail (1.58 < 5.05) with P = 0.31. Conversion factor ratio decreases and therefore the agreement in LP estimate between methods is closer during lower density nesting seasons.

**Fig 7.**
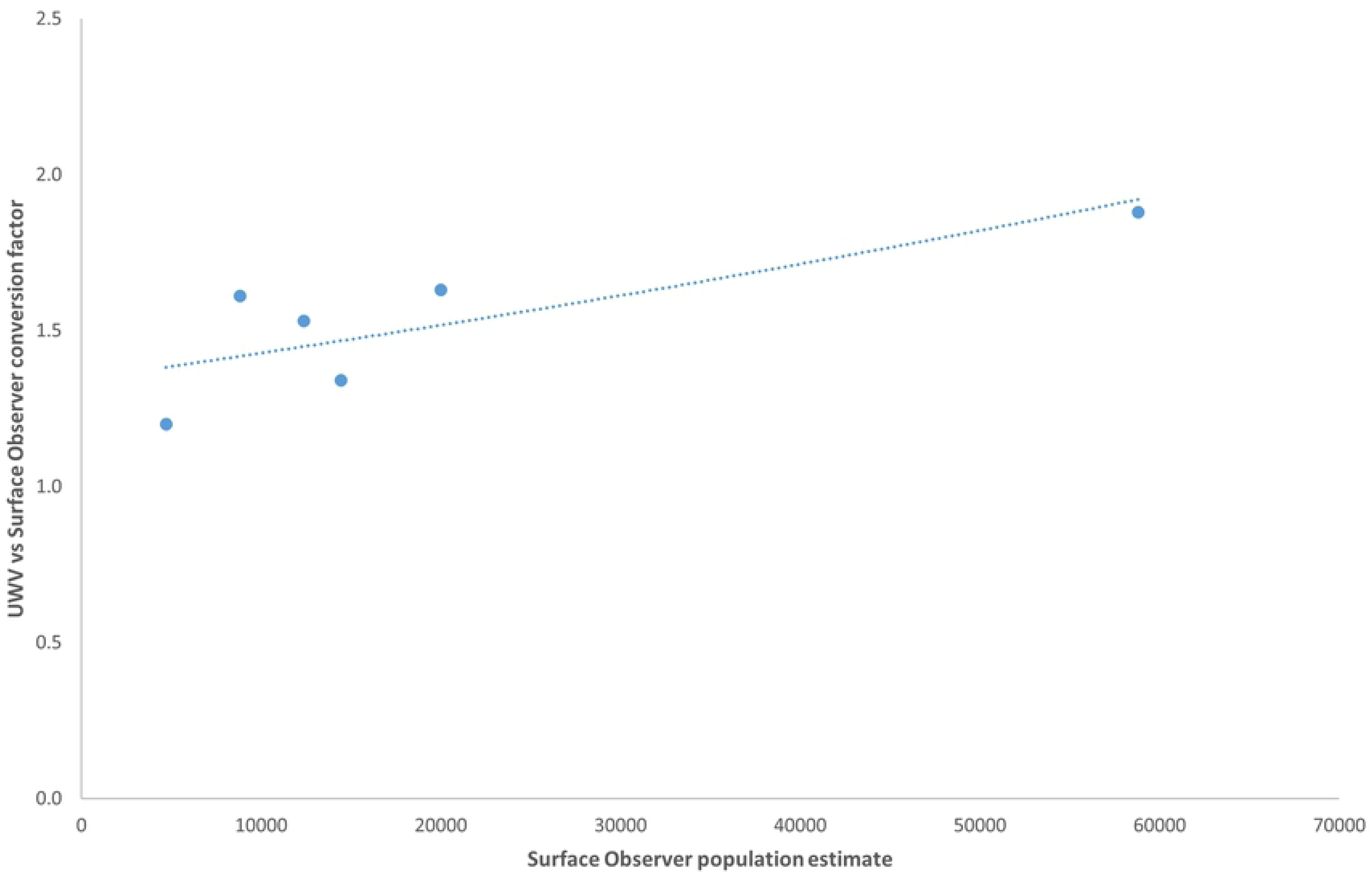
Conversion factor ratio of UVW vs SO methods compared with population estimates from SO surveys from different seasons. Fitted regression line y = 1.3436e^6E-06x^ has an *r^2^* value = 0.577

The use of UAVs to conduct mark/resight surveys is considerably more efficient in survey time (1:2.5 hrs) and personnel commitment (1:3) than the other survey techniques. UAV surveys can also be conducted in more extreme weather conditions (13:8 ms^−1^) while still providing precise estimates (Table 3). Consistent rain negates UAV flight options but is not a major impact on the other methods.

**Table 3.** Comparison of the cost effectiveness and logistical considerations for each turtle count method.

| Survey method | Equipment Cost | Personnel | Survey time | Viable wind conditions |
| --- | --- | --- | --- | --- |
| Surface observer | Low | 3 | 2.5 hrs | 8 ms<sup>-1</sup> |
| UWV | Low-moderate | 3 | 2.5 hrs | 8 ms<sup>-1</sup> |
| UAV | Moderate | 1 | 1 hr | 13 ms<sup>-1</sup> |

\*The survey time period refers to the total time to cover transects as shown on Figure 1.

UAVs also searched a larger search swath than the other two methods, resulting in 0.585 km^2^ searched on each occasion, compared to an estimated 0.4 km^2^ for the UWV method (assuming a distance of 10 m and a viewing angler of 127°) and 0.5 km^2^ for the SO method (assuming a search radius of 15 from the vessel). The average number of turtles counted by the UAV tended to be higher (3041) than the other methods (SO: 1228; UWV: 1345) (Table 2 and Fig 8). However, neither total numbers nor densities significantly differed between methods (log_e_ total numbers: D = 2.737, df = 2, p = 0.375; log_e_ density: D = 0.458; df=2, p= 0.795).

**Fig 8.**
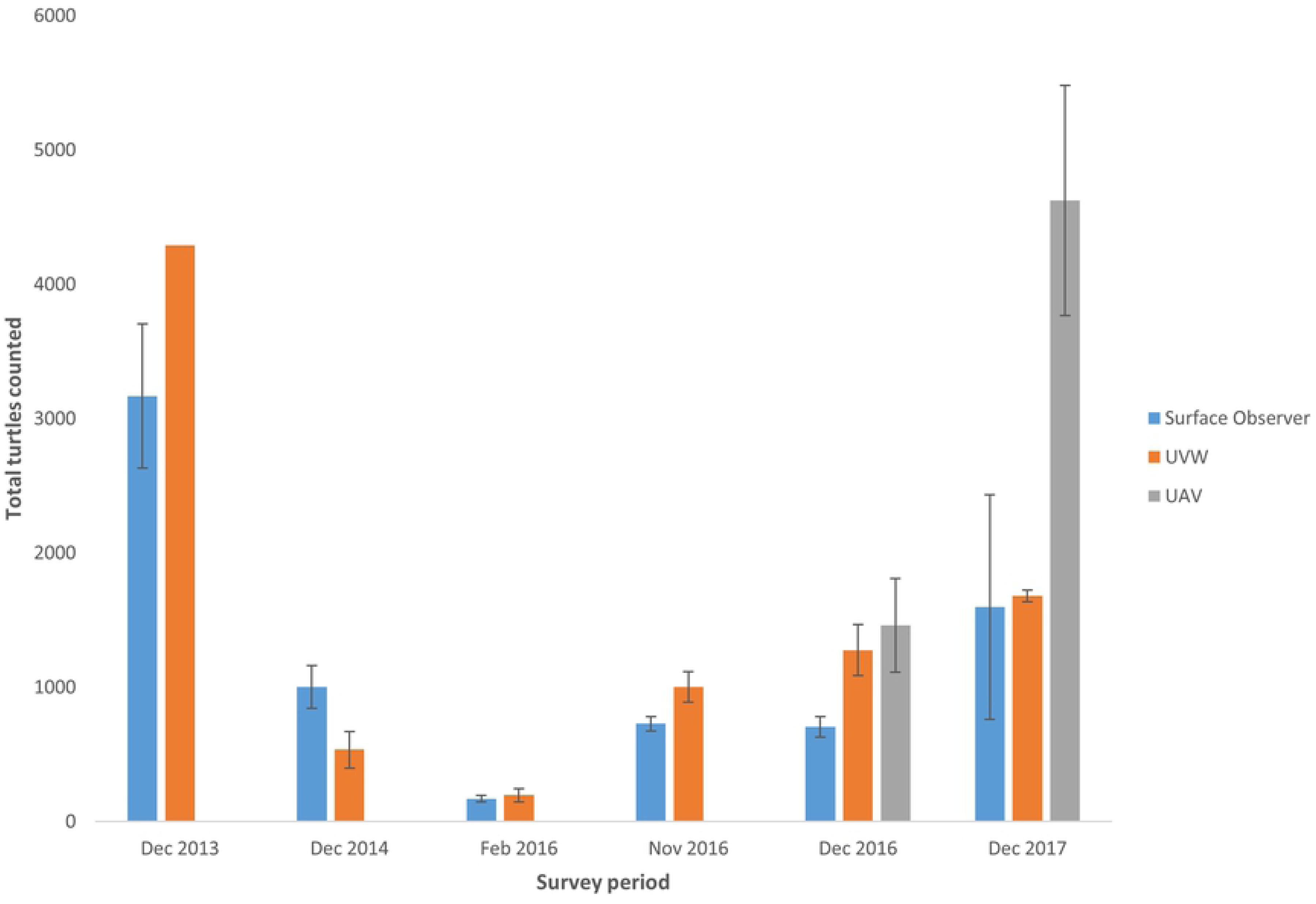
Mean total turtles counted (painted + unpainted) for periods surveyed by each method. Error bars shown are ± 1 standard error.

## Discussion

The UAV and UWV methods detected a lower ratio of marked to unmarked turtles than the SO method, resulting in considerably higher estimates of nester abundance. UAVs yielded an estimate 1.5x higher than the historical SO method, whereas the UWV method estimated 1.7x more turtles than the SO method. However, there was considerable variation in detection probabilities between sampling periods, which was likely to be driven by the extreme variability in the density of turtles in the inter-nesting habitat, suggesting that robust correction factors would require more sampling across a range of turtle densities.

A key advantage of the UWV and UAV approaches is the ability to review and playback video at speeds most suitable for accurate counts, especially when turtle aggregations were dense. The biased attraction of painted turtles to the observer’s eye is not tested or quantified but is considered to be the major factor causing the higher percentage of painted turtles recorded by the surface observer, resulting in lower overall population estimates by this method. We posit that the marked differences in the detection rate of marked turtles between the photographic and visual observer methods is due to visual searching limits of human observers. The performance of visual searching, as measured by search accuracy or reaction time, typically declines as the number of objects increases (Palmer 1994; Eckstein et al 2000). Observer fatigue can also influence detection rates (Lardner et al 2019). Further, in analysing complex natural or visually noisy scenes, humans direct visual attention towards regions of high contrast attract visual attention, particularly reflective surfaces such as white paint that represent high luminance contrast (Einhäuser et al 2003). This effect would be even greater in the noisy environment caused by surface reflections or surface disturbance (Lardner et al 2019). In our experiment, the white mark was discernible three meters deeper than the turtle model, suggesting that it may have drawn the attention of an observer who was subsequently able to discern that it was a turtle. Together, these mechanisms may explain why a visual observer had a higher probability of identifying marked turtles than the UAV or underwater video approach. This may also explain the fact that detection probability was the most similar between the underwater video and the visual observer in February 2016, when the population estimate was the lowest (Figs. 3&8 and Table 2). We predict that search accuracy would be greater when there are fewer turtles.

The in-water detectability of painted and unpainted turtles indicated that turtles were identifiable to 10 m depth, and that there were no pronounced differences in water clarity between sampling locations that were likely have influenced the results. However, we did not test how the viewing angle and surface conditions influenced detectability. Counting from the SO platform was mostly conducted at an angle to the surface of the water, and hence more subject to interference from glare and surface disturbance than the UAV or UWW method. This may have also influenced the ratio of painted to unpainted turtles detected, because the paint on remains visible during these conditions.

Compared to variation between sampling periods, there was little variation association with the timing of sampling or over consecutive samples. This suggests that the population is closed during sampling, an assumption also supported by the results of two other parallel studies. Firstly, the rate of mortality is low, with a maximum of 0.045% during the sampling period (interpolated from Robertson et al, in prep). Secondly, recently satellite tracking of 40 nesters at Raine Island in the 2017-18 and 2018-19 nesting seasons indicated that the vast majority of turtles remained in the immediate vicinity of the Raine Island reef edge after successful or unsuccessful laying. This study also supported a lack of bias in the location availability for detection of painted versus unpainted turtles. It demonstrated no significant difference between presence within the survey area during the first three days post nesting (the survey period) and the remaining internesting period (Mark Hamann, James Cook University, pers. comm.).

The use of UAVs to conduct mark-resight surveys is considerably more efficient in survey time (1:2.5 hrs) and personnel commitment (1:3) than the other survey techniques. Video analysis to count turtles is done manually at present however automated image analysis techniques are almost complete and will remove this extra time and personnel requirement. UAV surveys also still provide quality data when the sea-surface state and wind (i.e. 8-13 ms^−1^ winds) limit the SO or UWW methods, although consistent rain hinders the use of UAVs. The efficiency of the UAV method also facilitates cost-effective optimisation of the study design by using resampling to increase the precision of the population estimates (Fig 6).

The use of video recording versus the use of overlapping still images to produce a single orthomosaic image by UAV were both considered. For this application the benefit of moving video images during counting review provided the ability to adjust playback and pause footage to enable each individual turtle to be assessed as the UAV moved past. Movement was then used as part of authenticating turtle recognition, to gain different angle and reflectance aspects to optimise clarity of each turtle and paint mark and to allow the closest point of contact to be used in assessment (S2 Table).

Although no other studies have used UAVs in conjunction with mark-resight to estimate turtle abundance, other studies have used the direct count method, whereby counts of turtles are adjusted for the availability bias (Sykora-Bodie et al 2017). These adjustments were not deemed necessary in the Raine Island study due to the clear waters allowing detection to at least a 10 m depth range. The proportion of time spent by turtles in the detectable range to 10m depth is currently being investigated through studies of time depth recorders deployed on 21 nesters at Raine Island during the 2018-19 season. This will inform any bias of detectability for this mark-resight study and for use in total turtle counts conducted in other locations. Even acknowledging these limitations, our total and density estimates using the UAV survey method are higher than UAV density measurements of olive ridley turtles (*Lepidochelys olivacea*) at Costa Rica, the only other mass sea turtle nesting aggregation in the world. During the low-medium level nesting season in 2016 and the medium level nesting season in 2017 densities were 2496 ± 1441 turtles · km^−2^ and 7901 ± 1465 km^−2^ respectively. Low and high-end estimates of turtle density at Costa Rica were 1299 ± 458 km^−2^ and 2086 ± 803 km^−2^ respectively (Sykora-Bodie et al 2017).

## Conclusions

In summary, this study indicates that the use of UAVs for in-water mark-resight turtle population estimation is an efficient and accurate method that can provide an accurate adjustment for historical adult female population estimates at Raine Island. Underwater video may continue to be used as a backup method in case of UAV failure or weather restrictions to flight. This study also provides the basis for accurate nesting population estimation, including historical data correction, to inform reproductive success parameters for green turtles at Raine Island. This knowledge is crucial to identify the causes and quantify the levels of nesting and hatching failure and hatchling production. The data is also essential to the evaluation of improvements in reproductive success resulting from conservation management interventions such as re-profiling of the nesting beach and fencing to reduce adult female mortality (Dunstan, 2018).

## Acknowledgements

The Raine Island Recovery Project is a five-year, $7.95 million collaboration between BHP, the Queensland Government, the Great Barrier Reef Marine Park Authority, Wuthathi and Meriam Nation (Ugar, Mer, Erub) Traditional Owners and the Great Barrier Reef Foundation to protect and restore the island’s critical habitat to ensure the future of key marine species.

The funders had no role in study design, data collection and analysis, decision to publish, or preparation of the manuscript.

## Supporting information

**S1 Multimedia. UAV survey video.** Video of UAV survey at 50m altitude over waters adjacent to Raine Island reef edge.

**S1 Table.**
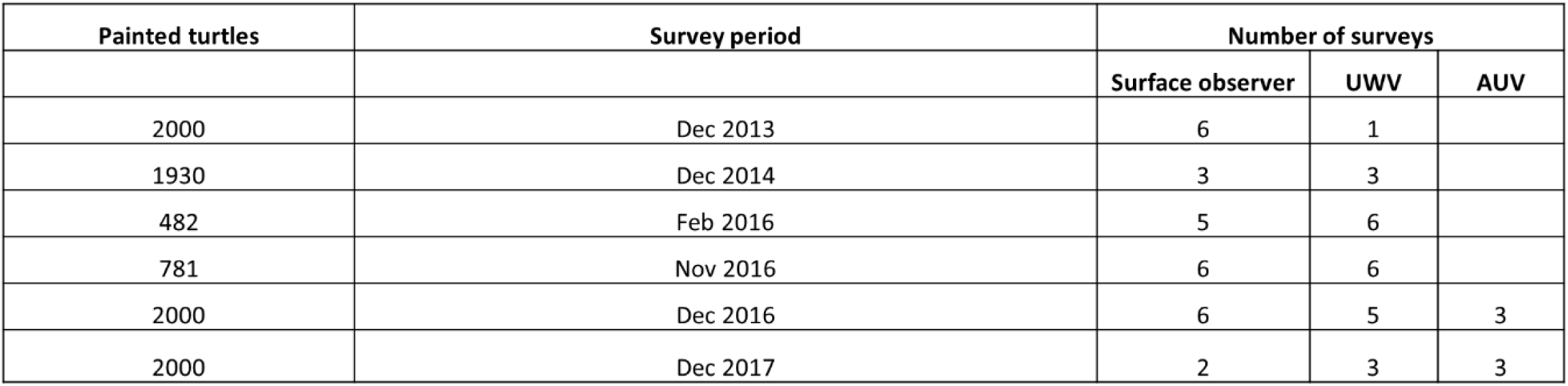
Survey area and estimated densities of turtles sighted using each method

| Painted turtles | Survey period | Number of surveys |  |  |
| --- | --- | --- | --- | --- |
|  |  | Surface observer | UWV | AUV |
| 2000 | Dec 2013 | 6 | 1 |  |
| 1930 | Dec 2014 | 3 | 3 |  |
| 482 | Feb 2016 | 5 | 6 |  |
| 781 | Nov 2016 | 6 | 6 |  |
| 2000 | Dec 2016 | 6 | 5 | 3 |
| 2000 | Dec 2017 | 2 | 3 | 3 |

